# Novel prosthecate bacteria from the candidate phylum Acetothermia revealed by culture-independent genomics and advanced microscopy

**DOI:** 10.1101/207811

**Authors:** Li-Ping Hao, Simon Jon McIlroy, Rasmus Hansen Kirkegaard, Søren Michael Karst, Warnakulasuriya Eustace Yrosh Fernando, Hüsnü Aslan, Rikke Meyer, Mads Albertsen, Per Halkjær Nielsen, Morten Simonsen Dueholm

## Abstract

Members of the candidate phylum Acetothermia are globally distributed and detected in various habitats. However, little is known about their physiology and ecological importance. In this study, an OTU belonging to Acetothermia was detected at high abundance in two full-scale anaerobic digesters. The first closed genome from this phylum was obtained by differential coverage binning of metagenomes and scaffolding with nanopore data. Genome annotation and metabolic reconstruction suggested an anaerobic chemoheterotrophic lifestyle in which the bacterium obtain energy and carbon via fermentation of peptides, amino acids, and simple sugars to acetate, formate, and hydrogen. The morphology was unusual and composed of a central rod-shaped cell with bipolar prosthecae as revealed by fluorescence *in situ* hybridization combined with confocal laser scanning microscopy, Raman microspectroscopy and atomic force microscopy. We hypothesize that these prosthecae allow for increased nutrient uptake by greatly expanding the cell surface area, providing a competitive advantage under nutrient-limited conditions.

## Main body

Life, as we know it, would not be possible without microbes. They catalyse key reactions in all of the major biogeochemical nutrient cycles^1^ and are being harnessed for their ability to convert waste products into renewable resources^2^. They are also essential for our health, both as beneficial members of our microbiome^3^ and as pathogens causing infections^4^. Knowledge of how distinct microorganisms fit into these processes is required to better predict the consequences of environmental changes and improve our health as well as the efficacy of biotechnological processes.

Culture-independent surveys of bacterial communities based on amplicon sequencing of 16S rRNA genes or concatenated single-copy phylogenetic marker genes have revolutionised our understanding of microbial community dynamics and diversity^2,5–7^ However, such analyses also reveal that many bacterial lineages lack cultivated representatives, and the bacteria affiliated to these candidate lineages are often poorly described^6,8,9^. These uncharted branches of the tree of life contain valuable information about the evolution of bacteria, exciting novel metabolic pathways, and hitherto unknown functions in microbial communities^6,10–13^.

The fast developments in next-generation sequencing and metagenomics enable the characterization of the whole community gene pool and can be used to elucidate the functional potential of individual microbial members. This allows us to better understand the ecological roles and interactions of the ubiquitous uncultivated microorganisms^14–17^. Genomes of uncultured microorganisms can be recovered from deeply sequenced metagenomes using different methodologies, such as the differential coverage binning approach^14^. Such attempts have been made to establish metabolic models and predict the ecophysiology of several candidate bacteria, such as *Candidatus* Fermentibacter daniensis (candidate phylum Hyd24-12)^18^, OP9/JS1 (candidate phylum Atribacteria)^19^ and *Candidatus* Promineofilum breve (phylum Chloroflexi)^20^.

In one of our studies of anaerobic sludge digesters^18^, a metagenome assembled genome (MAG) classified to the candidate phylum Acetothermia (former OP1)^8^ was found to be present in high abundance. The first draft MAG from this phylum was obtained from a subsurface microbial mat in the hot water stream^21^. It was predicted to possess a folate-dependent acetyl-CoA pathway of CO_2_ fixation and have an acetogenic lifestyle. Accordingly, it was given the name *Candidatus* Acetothermum autotrophicum. Another MAG (Acetothermia bacterium 64_32) was extracted from a marine shelf siliciclastic sandstone deposit from an oil reservoir^12^. This draft genome, however, lacked essential genes encoding for autotrophic CO_2_ fixation pathways, indicating a heterotrophic lifestyle. Other physiological information about this candidate phylum is currently not available.

Acetothermia bacteria occupy diverse habitats and have been detected in several geographically separated anaerobic digesters (**Figure S1**), suggesting that some members of this phylum may be specifically suited for this environmental niche and play a role in the conversion of organic matter into biogas. This motivated us to conduct a detailed investigation into the phylogeny, morphology, physiology, and ecology of Acetothermia bacteria in anaerobic digesters using amplicon sequencing, metagenomics, and advanced visualization techniques. This allows us, for the first time, to reveal an unusual morphology and physiology of this unrecognized microbial player in anaerobic digesters.

## Materials and methods

### Sample collection and storage

Between one and ten biomass samples were obtained from each of 31 anaerobic digesters treating primary and surplus sludge at 18 Danish wastewater treatment plants (WWTPs) in the period from 2011 to 2017 (**Table S1, S2**). A volume of 50 mL digester biomass was sampled, homogenized, and stored as 2 mL aliquots at −80°C before DNA extraction. DNA was extracted using the FastDNA Spin kit for soil (MP Biomedicals, Santa Ana, CA, USA) as optimized for anaerobic digesters by Kirkegaard *et al.*^18^.

### Amplicon sequencing of the 16S rRNA gene

The V4 variable region of the bacterial and archaeal 16S rRNA gene was amplified with the PCR primers 515F^22^ (3’-GTGCCAGCMGCCGCGGTAA) and m806R (3’-GGACTACNVGGGTWTCTAAT) and sequenced using the Illumina platform as described by Albertsen *et al.^23^.* The m806R primer is a modified version of 806R^22^, in which the degeneracy of a single base is increased to ensure a perfect match to all Acetothermia sequences in the SILVA SSU Ref NR 99 database (Release 128)^24^.

Amplicon sequencing libraries (**Table S2**) were pooled in equimolar concentrations with a final loading concentration of 10 pM and sequenced on the MiSeq (Illumina) platform using a MiSeq reagent kit v3 (2 × 300 bp).

All sequenced sample libraries were trimmed, and low quality reads were removed using trimmomatic v. 0.32^25^ and then merged using FLASH v. 1.2.11^26^. The reads were screened for potential PhiX contamination using USEARCH v. 7.0.1090^27^. The reads were clustered into operational taxonomic units (OTUs, sequence identity ≥ 97 %) using USEARCH and subsequently classified using the RDP classifier^28^ with the MiDAS database v. 1.23^2^. Further analyses were performed in R environment v. 3.4.1^29^ using the R CRAN packages ampvis v. 1.24^23^ and ggplot2 v. 1.0.1^30^. The samples were subsampled to an even depth of 10,000 reads per sample.

### Illumina sequencing, metagenome assembly, and genome binning

Illumina Nextera DNA Library Prep kit was used to prepare metagenome libraries following the standard protocol. DNA extracts from four samples collected at different time points during the first half of 2016 from a mesophilic digester at Randers WWTP were used as templates for library preparation (**Table S3**). The libraries were paired-end (2 × 250 bp) sequenced on the Illumina HiSeq 2500 platform using the HiSeq Rapid PE cluster kit v2 and the HiSeq Rapid SBS kit v2 (500 cycles) in rapid run mode and also paired-end sequenced (2 × 300 bp) on Illumina MiSeq platform using MiSeq reagent v3 (600 cycles). Standard protocols were used for sample preparation and sequencing. The metagenomic assembly and binning process was performed as described by Kirkegaard *et al.*^18^.

### Nanopore sequencing

Genomic DNA was prepared for 1D nanopore sequencing (Oxford Nanopore Technologies, UK), following the manufacturer’s protocol (LSK-108) without the optional DNA shearing and DNA repair steps. The library was loaded on a FLO-MIN106 flow cell and sequenced using the MinION Mk1B DNA sequencer (Oxford Nanopore Technologies). The sequencing software used was MinKNOW v. 1.7.3 (Oxford Nanopore Technologies, UK) with the 48-hour sequencing workflow (NC_48Hr_Sequencing_Run_FLO_MIN106_SQK-LSK108.py). Sequencing reads were base-called using Albacore v. 1.2.1 (Oxford Nanopore Technologies, UK).

### Genome closing and annotation

The SSPACE-LongRead scaffolder v. 1.1^31^ was used to assemble contigs from the Acetothermia genome bin into a single scaffold based on the long Nanopore reads. Gaps in the draft genome were closed using GapFiller v. 1.11^32^ or by manual read mapping and extension in CLC Genomics Workbench v. 9.5.2. Finally, the closed genome was manually polished to remove SNPs and ensure a high quality assembly (**Table S4**). Genome annotation was performed in the ‘MicroScope’ annotation pipeline^33,34^ as described by Kirkegaard *et al.*^18^.

### Phylogeny of the 16S rRNA gene and FISH probe design

Phylogenetic analysis and FISH probe design were performed using the ARB software package^35^ with the SILVA SSURef NR 99 database (Release 128)^24^. All sequences classified to the Acetothermia phylum from the SSURef database were included, except those from the same study that shared ≥ 99% similarity. Potential probes were assessed *in silico* with the mathFISH software^36^. The Ribosomal Database Project (RDP) PROBE MATCH function^37^ was used to identify non-target sequences with indels^38^. Probe validation and optimization were based on generated formamide dissociation curves, as described by Daims *et al.*^9^. The final probes are shown in **Table S5** and have been deposited in the probeBase database^40^.

### Sample fixation and fluorescence *in situ* hybridization (FISH)

Fresh biomass samples, taken from sludge digesters at Randers and Esbjerg WWTPs, were treated by either ethanol or paraformaldehyde (PFA) for the optimal fixation of Gram-positive and Gram-negative bacteria, respectively. For PFA fixation, diluted samples [1:4 in 1 × Phosphate-Buffered Saline (PBS) solution] were first fixed with 4% (w/v) PFA and then stored in 50% (v/v) ethanol / 1 × PBS solution at −20°C, as previously described^39^. For ethanol fixation, pellets were first obtained by removing the supernatant by centrifugation at 12,000 g for 5 min at 4°C, and then directly fixed and stored in 50% (v/v) ethanol / 1 × PBS solution.

FISH was performed as detailed by Daims *et al.^39^.* The hybridization conditions applied for each probe are given in **Table S5**. The NON-EUB probe was applied as a negative control for hybridization^41^. Nucleic acids in cells were stained with either 4’,6-diamidino-2-phenylindole (DAPI) (50 μM, for 30 min) or Syto9 (6 μM, for 20 min) (Molecular Probes, Eugene, Oregon, USA). Microscopic analysis was performed with an Axioskop epifluorescence microscope (Carl Zeiss, Oberkochen, Germany) or a white light laser confocal microscope (Leica TCS SP8 X) fitted with a 405 nm diode laser (Leica Microsystems, Kista, Sweden). Excitation (Ex.) and emission (Em.) details applied are as follows: DAPI (Ex. 405 nm; Em. 440-615 nm); Syto9 (Ex. 485 nm; Em. 490-550 nm); FLUOS (Ex. 492 nm; Em. 500-555 nm); Cy3 (Ex. 554 nm; Em. 565-650 nm); Cy5 (Ex. 649 nm; Em. 660-695 nm).

### Raman spectroscopy

To locate the Acetothermia cells for Raman analysis, FISH was conducted on optically polished CaF_2_ Raman windows (Crystran, UK) by using the newly designed OP1 probes labelled with Cy3. Once the cells were located, the fluorescence of Cy3 was bleached by keeping the Raman laser on the target cell for 5 min. Raman spectra from single cells of Acetothermia were obtained using a Horiba LabRam HR 800 Evolution (Jobin Yvon-France) equipped with a Torus MPC 3000 (UK) 532 nm 341 mW solid-state semiconductor laser. Prior to all measurements, the Raman microspectrometer was calibrated to the first order Raman signal of Silicon occurring at 520.7 cm^−1^. The CaF_2_ Raman substrate also contains a single sharp Raman marker at 321 cm^−1^, which serves as an internal reference point in every spectrum. The incident laser power density on the sample was attenuated down to 2.1 mW/μm^2^ using a set of neutral density (ND) filters.

The Raman system is equipped with an in-built Olympus (model BX-41) fluorescence microscope. A 50x, 0.75 numerical aperture dry objective (Olympus M Plan Achromat-Japan), with a working distance of 0.38 mm was used throughout the work. A diffraction grating of 600 mm/groove was used, and the Raman spectra collected spanned the wavenumber region of 200 cm^−1^ to 1800 cm^−1^. The slit width of the Raman spectrometer and the confocal pinhole diameter were set to 100 pμm and 150 μm, respectively. Raman microspectrometer operation and subsequent processing of spectra were conducted using LabSpec version 6.4 software (Horiba Scientific, France).

### Atomic force microscopy (AFM)

The combined optical and atomic force microscopy experiments were carried out with a sample stained with 10 μM Syto9 in PBS solution. A JPK Nanowizard IV system (Berlin, Germany) on an inverted Zeiss Axiovert 200M epifluorescence microscope was used, with a 63x oil immersion optical lens (Zess Plan-Apochromat, NA 1.4) and Zeiss filter set 10 (Ex. 450-490 nm, Em. 525-565 nm). This AFM setup was used in the QI™ mode, which is a dynamic nanomechanical mapping (DNM) method. This DNM method can simultaneously provide height channels for morphology, and force spectroscopy based information, i.e. adhesion channel and force-distance curves. Although DNM methods are often used for accessing mechanical and physicochemical properties of the sample, they are also employed for high-resolution imaging due to their capability to directly control the tip-sample interaction forces below nanoNewton level. In this work, DNM was employed for advanced imaging, and the scans were acquired with a soft cantilever, namely Scanasyst-Air (Bruker). Nominal values for the cantilever’s resonance frequency and the spring constant are 70 kHz and 0.4 N/m, respectively. The operation parameters such as set point and Z length were varied to optimize the scan for the highest resolution and to minimize the risk of damaging the tip and the sample. The pixel time was kept at 30 ms. Z range was set to 15 μm, and the images were initially acquired with 256×256 px, then with 512×512 px, if possible. All DNM experiments were carried out in air under room conditions.

### Data availability

All sequencing data has been submitted to the ENA under the project ID PRJEB22104. Amplicon sequencing data is available with the accession numbers ERS1910092-ERS1910183, metagenome data with the accession numbers ERS1909451-ERS1909457, and the complete genome under accession number ERZ478283.

## Results and discussion

Acetothermia bacteria have previously been observed in anaerobic digesters^42^, but their distribution and abundance in these systems are not known. It was therefore decided to survey the microbial composition of 31 full-scale anaerobic digesters using 16S rRNA gene amplicon sequencing. The choice of PCR primers can have a pronounced bias on the microbial composition observed^23^. Three common 16S rRNA gene amplicon primer pairs were evaluated on samples from a digester containing Acetothermia, and metagenomes of the same samples derived from primer-independent shotgun sequencing were used as references (**Figure S2**). Only the 515F/806R^22^ primer pair, which targets V4 region of the 16S rRNA gene, was able to amplify Acetothermia-related 16S rRNA genes and provide an estimate of Acetothermia relative abundance in the samples. To ensure that the primer pair would be able to target all Acetothermia, we compared the primer sequences to Acetothermia sequences in the SILVA databases. It was found that 67.4% of the sequences contained a single mismatch to the 806R primer. However, this could be alleviated by increasing the degeneracy of the primer at a single position. This modification did not affect the overall community structure of the samples tested, so the modified primer was used for the survey.

Only a single genus-level OTU assigned to phylum Acetothermia was observed in four mesophilic sludge digesters at two WWTPs from the survey of 31 digesters (**Figure 1A**). The OTU was stably present over a period of three to six years in these digesters, but displayed a notable decline from the summer of 2016. It ranked among the five most abundant bacterial OTUs and constituted from 0.1 to 8.9% of all sequenced 16S rRNA gene amplicons (**Figure 1B**). The Acetothermia OTU was not detected in amplicons of the incoming feed streams (primary and surplus biological sludge from the wastewater treatment processes), which indicates that the abundance observed was due to growth in the digesters and not immigration. No OTUs related to Acetothermia were observed in thermophilic digesters, or the mesophilic digester operated with thermal hydrolysis of feedstock **(Figure 1A**). This indicates that the Acetothermia OTU has special habitat requirements specific to some mesophilic systems that treat primary and surplus biological sludge.

**Figure 1.**
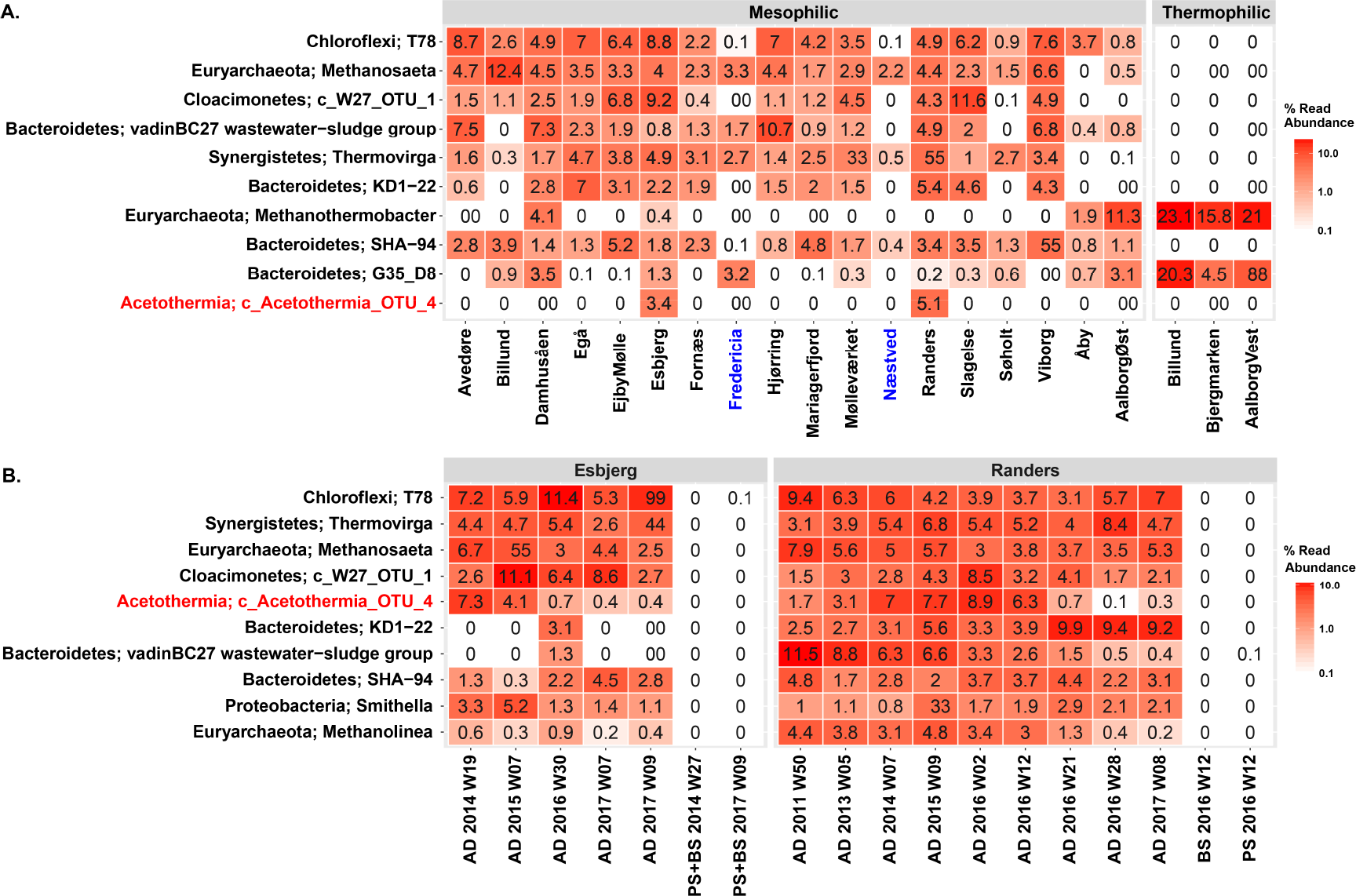
Heatmap of the ten most abundant microbial genera in anaerobic digesters treating sewage sludge. (A) Average genera abundances of the period of 2011~2016 in the digesters from 20 wastewater treatment plants (WWTPs). Labels at the bottom of the heatmap represent the location of WWTPs and digesters. Blue labels represent WWTPs applying thermal-hydrolysis process for pre-treating the feedstock. (B) Temporal analysis of the microbiome composition in the digesters from Randers and Esbjerg WWTPs and of the feedstock. Mean abundances of two digesters running in parallel at each WWTP were shown in the profile. Labels at the bottom of the heatmap represent sample type, year, and week of sampling time. Sample type includes: AD for sludge from anaerobic digesters; PS for sludge from the primary clarifier, and BS for surplus biological sludge from secondary clarifier; BS+PS for a mixture of PS and BS before being fed into the digester. Classification levels presented are phylum and genus, which are separated by semicolon. The genera are sorted by the mean abundance across all the analyzed samples.

### Complete genome of the Acetothermia bacterium

To learn more about the ecophysiology of Acetothermia bacteria in anaerobic digesters, we sought to obtain genomic information from the abundant OTU. This organism was consistently found in high abundance in a full-scale anaerobic sludge digester at Randers WWTP, thus providing a good system for in-depth investigations (**Figure 1B**). To this end, metagenomes were constructed from four individual biomass samples collected during the first half of 2016 (**Table S3**) and a 12-contig draft genome of Acetothermia bacterium sp. Ran1 (‘Ran1’ in short) was successfully binned from these using differential coverage binning^14^ (**Figure S3**). Long read Nanopore data was obtained from one of the four samples and used to scaffold the draft genome and create a complete closed genome after manual polishing (**Table S4**).

The closed genome was 1.32 Mbp and had a GC content of 68.2%. The genome encodes a single split rRNA operon, in which the 16S rRNA gene was located away from the co-localized 23S and 5S rRNA genes. Fewer rRNA gene copies, as well as more split rRNA operons, are common features in host-dependent bacteria, but less frequent in free-living cells^43^. Accordingly, Ran1 could be a host-dependent organism, but microscopy revealed that this was not the case for Ran1 (see below). Instead, we hypothesize that the anaerobic digester may provide a stable environmental niche for the bacterium, similar to that provided by a host cell.

### Phylogenetic analyses of Ran1

Ran1 was classified to the Acetothermia based on its 16S rRNA gene using the SILVA taxonomy. Phylogenetic analyses of the available sequences for this phylum revealed evident separation of lineages with similar ecological preferences and habitats (Figure 2A, **Figure S4**). The 16S rRNA gene sequence of Ran1 clustered into a mono-phylogenetic group together with sequences from other anaerobic digesters^42,44–47^. Based on the recommended sequence similarity cut-off values for the definition of phylogenetic taxa^48^, this group represents a new genus, within the same family as the uncultured Acetothermia bacterium 64_32^12^. A phylogenetic tree based on concatenated single copy marker gene was created and used to establish a broader phylogenetic context (**Figure 3**). This revealed that Ran1 and the previous Acetothermia draft genomes^12,21^ are distantly related to all currently available genomes, supporting its status as a novel phylum.

**Figure 2.**
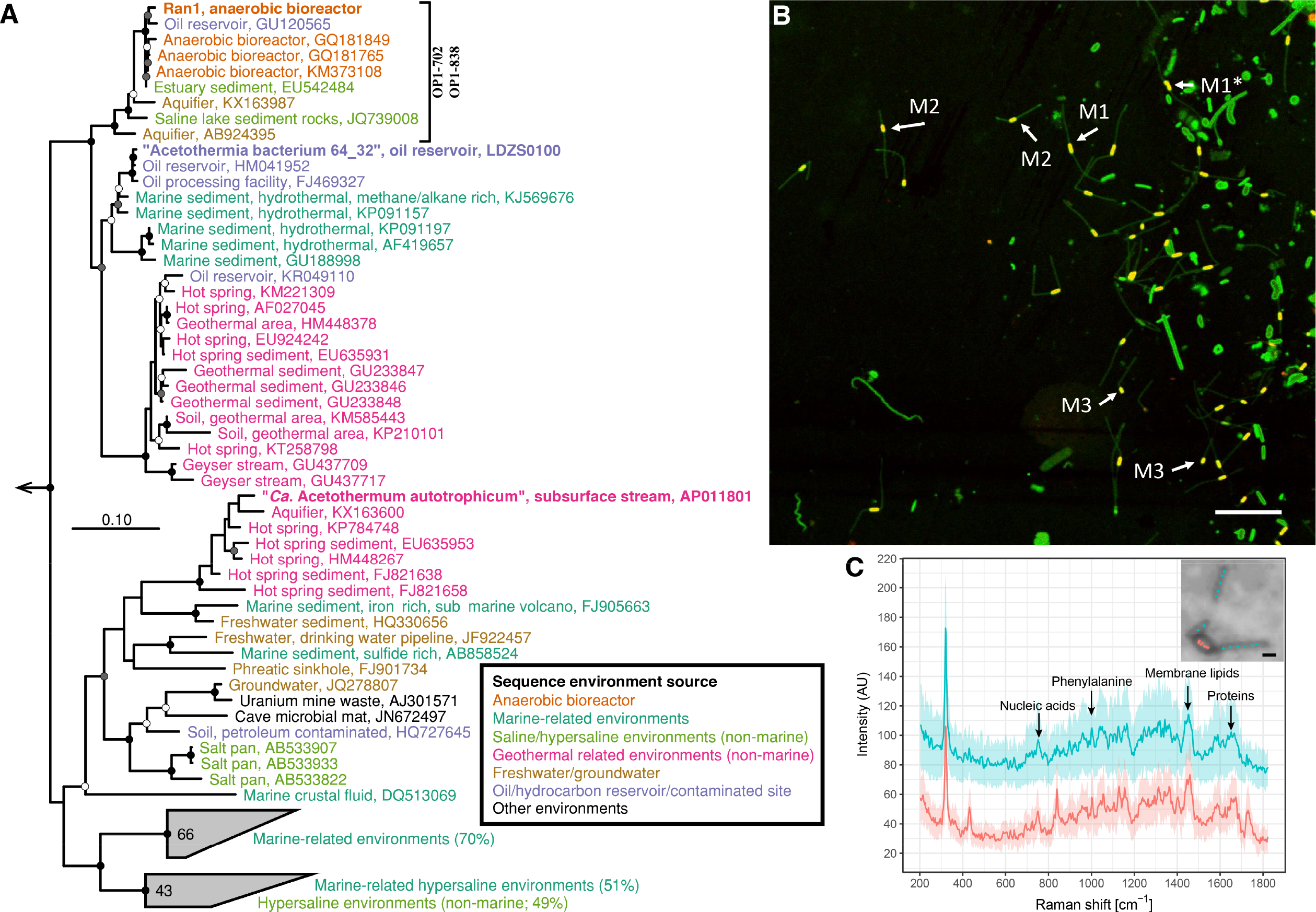
(A) Maximum-likelihood (PhyML) 16S rRNA gene phylogenetic tree of sequences classified to the candidate phylum Acetothermia (SILVA SSURef NR 99, Release 128). The alignment used for the tree applied a 20% conservational filter to remove hypervariable positions, giving 1120 aligned positions. Sequences are colored according to their isolation source environment. Proposed phylogenetic classification of the novel genus and coverage of the newly-designed FISH probes are indicated with a black bracket. Bootstrap values from 100 re-samplings are shown for branches with >50% (white dot), 50~70% (grey) and >90% (black) support. Species of the phylum Thermotogae were used as the outgroup. The scale bar represents substitutions per nucleotide base. An expanded version of the tree is provided in **Figure S4**. (B) Composite fluorescence micrograph of the Acetothermia cells, hybridized with the 0P1-702 FISH probe (Cy3, red) and stained with Syto9 (green). Yellow signal indicates overlap of fluorescence from Cy3 and Syto 9.Arrows indicate three slightly different morphologies: M1 = central rod with bipolar prosthecae of similar length; M2 = smaller central rod with bipolar prosthecae of different lengths; M3 = smallest central rod with a single polar prostheca. An M1 cell which seems to be undergoing cell division is indicated with an asterisk. PFA-fixed biomass samples were used, originating from an anaerobic sludge digester at Randers WWTP. Scale bar represents 10 μm. (C) Raman spectra of a bipolar prosthecate cell targeted by 0P1-702 probe. Seven spectra for the main rod body (red) and thirteen for the prosthecae (cyan) were obtained as indicated by the spots on the embedded cell image. Average spectra of the rod and prosthecae, respectively, are shown with the standard deviation depicted as ribbons. Peaks were assigned for nucleic acids (784 cm^−1^), phenylalanine (1004 cm^−1^), membrane lipids (1450 cm^−1^), and amide I linkages of proteins (1660 cm^−1^)^80^,^81^.

**Figure 3.**
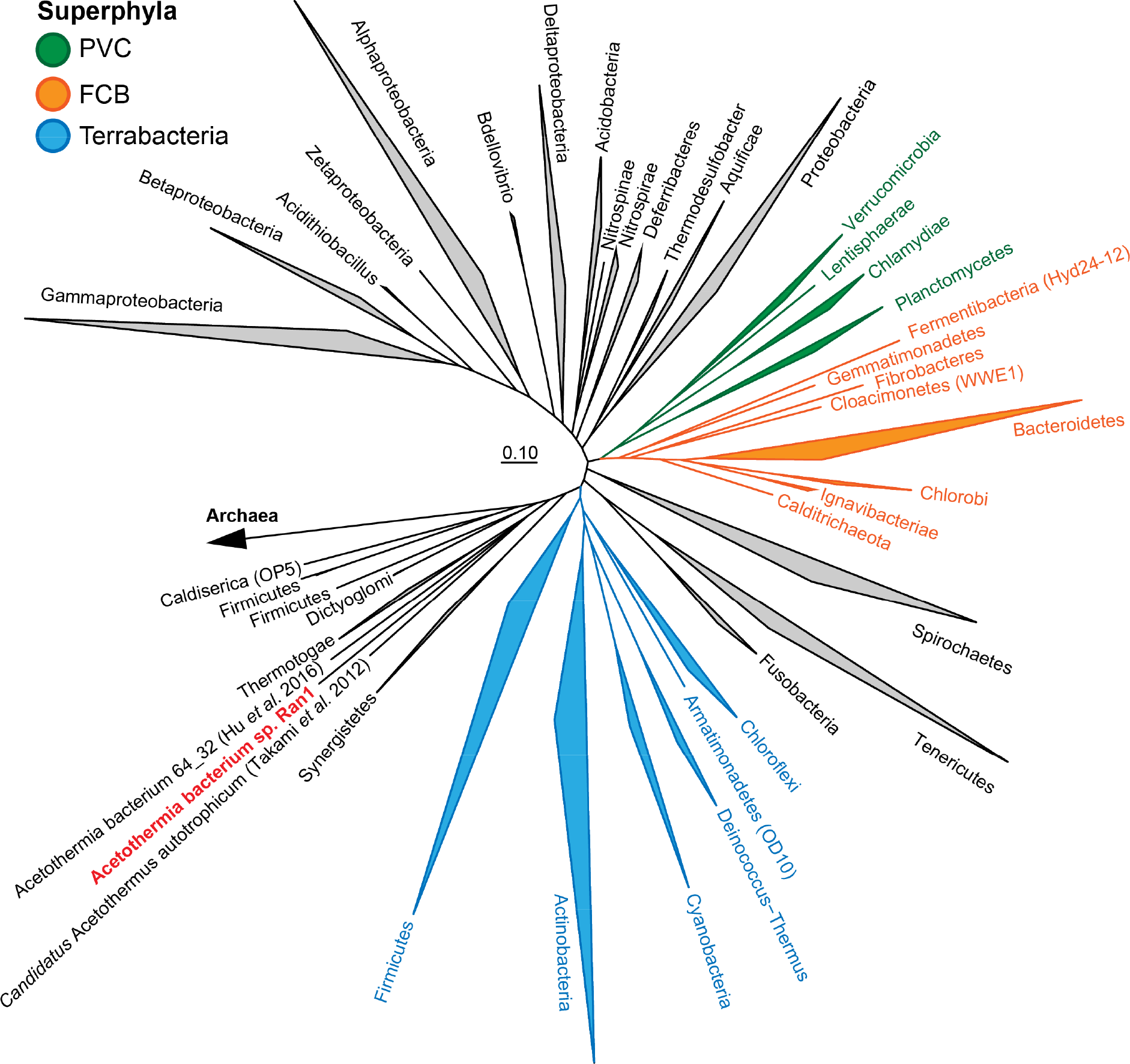
Phylogenetic position of Acetothermia genomes in the reference genome tree generated by CheckM v. 1.0.6^83^ and visualized in ARB v. 6.0.2^35^. The CheckM tree is inferred from the concatenation of 43 conserved marker genes and incorporates 2052 finished and 3605 draft genomes from the IMG database^83^.

### Morphology

To investigate the morphology of Ran1, we designed two FISH probes that cover the proposed novel genus that contains Acetothermia bacteria associated with anaerobic digesters (**Figure 2A**). These probes were then applied to samples from one of the digesters at Randers WWTP (Figures 2B, **S5, and S6**). Both PFA- and ethanol-fixed samples were analyzed to ensure optimal fixation of Gram-positive and Gram-negative bacteria, respectively (**Figure S6**). FISH results revealed small rod-shaped cells (approx. 0.8 x 1~2 μm) dispersed in the liquid phase, which were hybridized with the genus-specific probes. With ethanol-fixed biomass, appendages (approx. 0.4 x 4~8 μm) were observed at both poles of the rod-shaped cell. FISH signals for these structures were patchy, indicating a relatively low number of ribosomes present inside the appendages. No FISH signal were observed for the appendages with PFA fixed cells (**Figure S6**). When using Syto9 to stain the nucleic acids, these appendages were clearly visualized for the probe-hybridized cells in both PFA-and ethanol-fixed samples (**Figure S6**). It suggests that the nucleic acid containing cytoplasm was shared between the rod-shaped “main body” and the appendages. This was further confirmed by Raman microspectroscopy analysis, which demonstrated a similar composition in terms of nucleic acids, membrane lipids, and proteins of the main body and the appendages (**Figure 2C**). Probe-targeted cells from another digester at Esbjerg WWTP demonstrated similar morphology. Accordingly, we hypothesize that the appendages are extensions of the cell envelope out of the central rod body, similar to the prosthecae of *Caulobacter* and *Asticcacaulis*^49^.

Further analysis of FISH data demonstrated three different morphologies according to the size of the central rod and the length or appearance of the polar prosthecae: 1) central rod with bipolar prosthecae of similar length; 2) smaller central rod with bipolar stalks of different length; 3) smallest central rod with a single polar prostheca (**Figure 2B**). These different morphologies likely represent sequential development of bacterial morphology at different growth stages, in which small rods with a single prostheca represent cells just after cell division, and the longer rods with two prosthecae of equal length represent cells just before cell division. Indeed, it was possible to identify a few dividing cells with prosthecae of equal length (**Figure 2B**). Dynamic morphology change in a cell cycle is already known from other prosthecate bacteria, such as *Caulobacter*^53^ and *Asticcacaulis*^54^.

Higher resolution information on cell surface properties of Ran1 was obtained using atomic force microscopy (AFM) (**Figure 4**). AFM confirmed the morphology observed by FISH microscopy, i.e. a central rod-shaped cell with prosthecae extending from both poles. Analysis of four individual Ran1 cells revealed that the average width and length of the main rod body were 0.46±0.03 μm and 1.58±0.39 μm, respectively. The average height of only 0.066±0.017 μm showed that cells collapsed during air drying of the sample. The width of the prosthecae was relatively constant (0.256±0.004 μm), but decreased to 0.225±0.001 μm in cross sections where bending of the prosthecae occured. Such bendings were observed in most samples, and the degree of narrowing varied, based on the bending angle, which was up to 124.2±3.6°. This indicated flexibility of the prosthecae. The total length of the bacteria with prosthecae was 11.42±1.49 μm.

**Figure 4.**
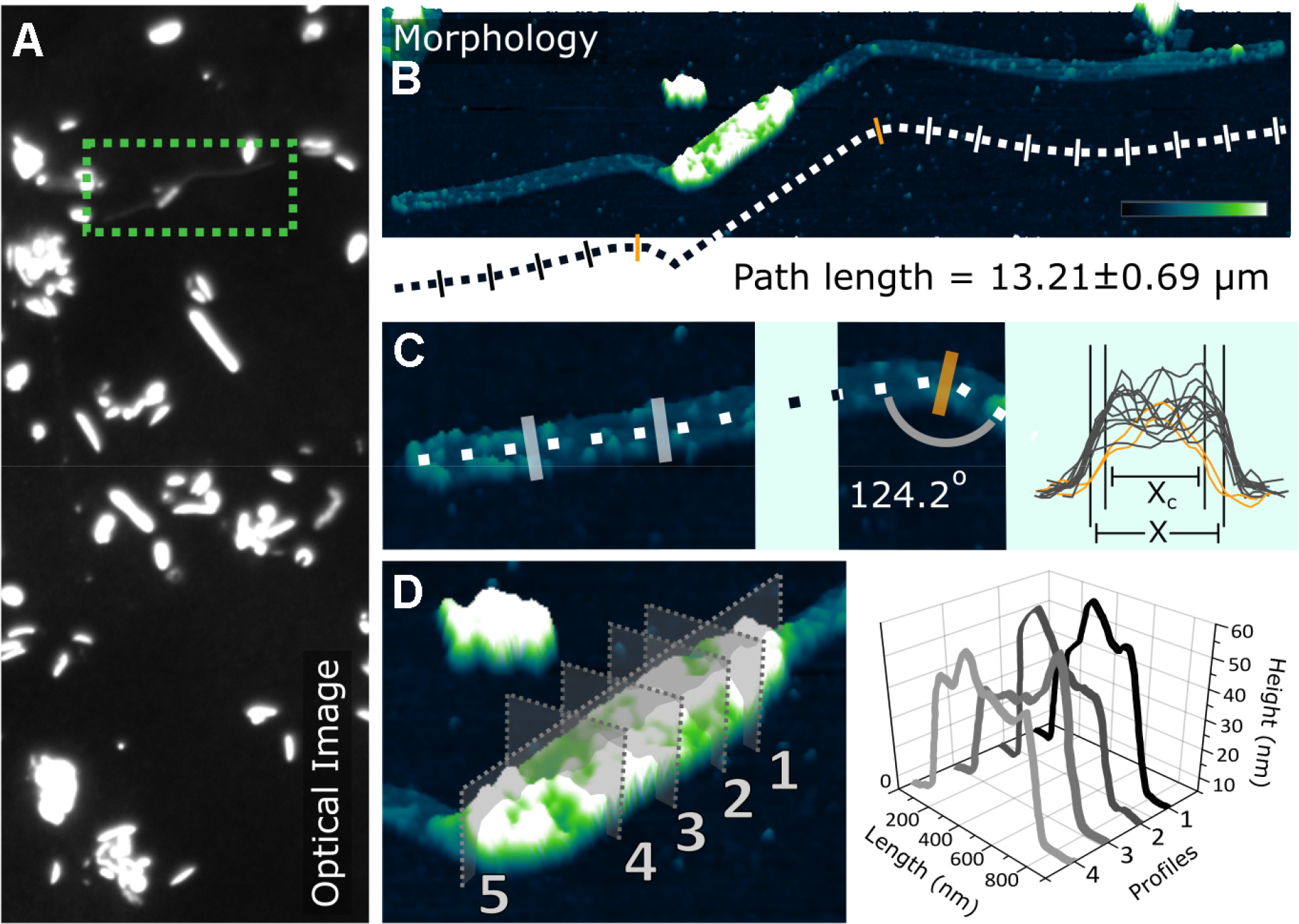
Combined optical and atomic force microscopy images reveal the morphology of Rani cells. (A) The optical image to the left shows a broad overview of the sample which is composed of bacteria of different shapes; (B) The morphology image presents the 3D form of a Ran1 cell in real space. The scale bar is 2 (xm in length, and the color transition represents the height change from 0 to 39 nm. (C) The cell stretches out to 13.21±0.6 μm with prosthecae at both poles, which are 0.26053±0.00911 μm (X) in width, except for slight narrowing down to 0.22465±0.00115 μm (Xc) due to bending with angles of up to 124.2±3.6°; (D) Zoom in the image of the main rod body. Cross sections show a rugged surface, as depicted and measured by Profile 1-4 perpendicular to the length of the rod represented by Profile 5.

The total surface area (SA) and surface area to volume ratio (SA/V) were calculated for the rod-shaped cell with and without the prosthecae, based on the observed average length and width. Results show that development of the prosthecae made the surface area increase by 3.5 times (from 2.28 to 10.20 (μm^2^) and SA/V become 42% larger (from 9.6 to 13.7 μm^−1^), providing an increased interface for nutrients uptake^50–52^. It has been demonstrated that prosthecate bacteria have a competitive advantage under nutrient deficient conditions, and they are often observed under such conditions^51-54^. Nevertheless, this effect is even more pronounced in diffusion-limited environments, where the rate of nutrient uptake is proportional to the effective linear dimension of a structure, rather than to its surface area^51^. Indeed, the length of the prostheca of *Caulobacter* inversely correlates with the availability of phosphate, indicating enhanced phosphate uptake capability^55^. Consistent with this observation, the digesters which harbour Ran1 in abundance demonstrated relatively low soluble orthophosphate concentration (around 25~80 mg PO_4_−P/L), compared to the other digesters (95~480 mg PO_4_−P/L) (**Table S1**). Furthermore, it was observed that the decrease of Ran1 (from 6~8% to < 1%) in the summer of 2016 followed an increase of phosphorus content (PO_4_−P and Total P) as well as concentration of organic compounds (VFAs, CODs) in the liquid phase (**Figure S7 and S8**). This supports the idea that Ran1 may have a competitive advantage in nutrient-limited engineered systems, especially at levels with relatively low amounts of phosphorus. Ran1 may, therefore, be used as a bioindicator for such a condition, but more studies are needed to verify this hypothesis.

### Genome inferred surface properties

Some cell envelope properties can be inferred directly from genomes, based on the presence or absence of cell envelope genes found specifically in archetypical mono- or diderm lineages^14^. This study revealed an unusual cell envelope architecture of Ran1, with similarities to both members of the monoderm Chloroflexi and the atypical diderms Thermotoga and Deinococcus-Thermus (**Figure S9**). Accordingly, it is less than easy to conclude whether Ran1 has a mono- or diderm cell envelope. The genome did not contain any genes associated with lipopolysaccharides, which are commonly found in the outer membrane of diderm bacteria^56^. However, genes encoding an outer-membrane-specific bacterial surface antigen and an outer-membrane permease imply that Ran1 may have a simple diderm cell envelope similar to those found in Thermotoga^57^. The sheath-like outer membrane of Thermotoga changes its size according to environmental conditions, which has been proposed to provide increased access to nutrients in the same manner as the prosthecae of prothecate bacteria^58^. Accordingly, it may be proposed that the outer membrane of Ran1 is a simple scaffold for high affinity nutrient transporter^51^.

Further genome annotation and specialized searches using the PilFind program^59^ did not reveal any genes associated to flagella, fimbriae, pili, or cell surface adhesins. However, a few genes related specifically to prostheca development were encoded by the genome, such as the bactofilin family cytoskeletal protein CcmA and the bifunctional penicillin-binding protein Pbp^53^. In *Caulobacter crescentus*, bactofilins are found as membrane-associated clusters at the pole of the cell, where they recruit the peptidoglycan synthase PbpC and initiate prosthecae development^53^. It is, therefore, likely that Ran1 may use a similar strategy for this purpose. Metabolic model for Ran1

To learn more about the potential function of Ran1, we constructed a metabolic model based on the annotated genome (**Figure 5** and **Table S6**). A brief overview of the metabolic model is provided below, and detailed descriptions of selected pathways are given in **Supplementary Results**.

**Figure 5.**
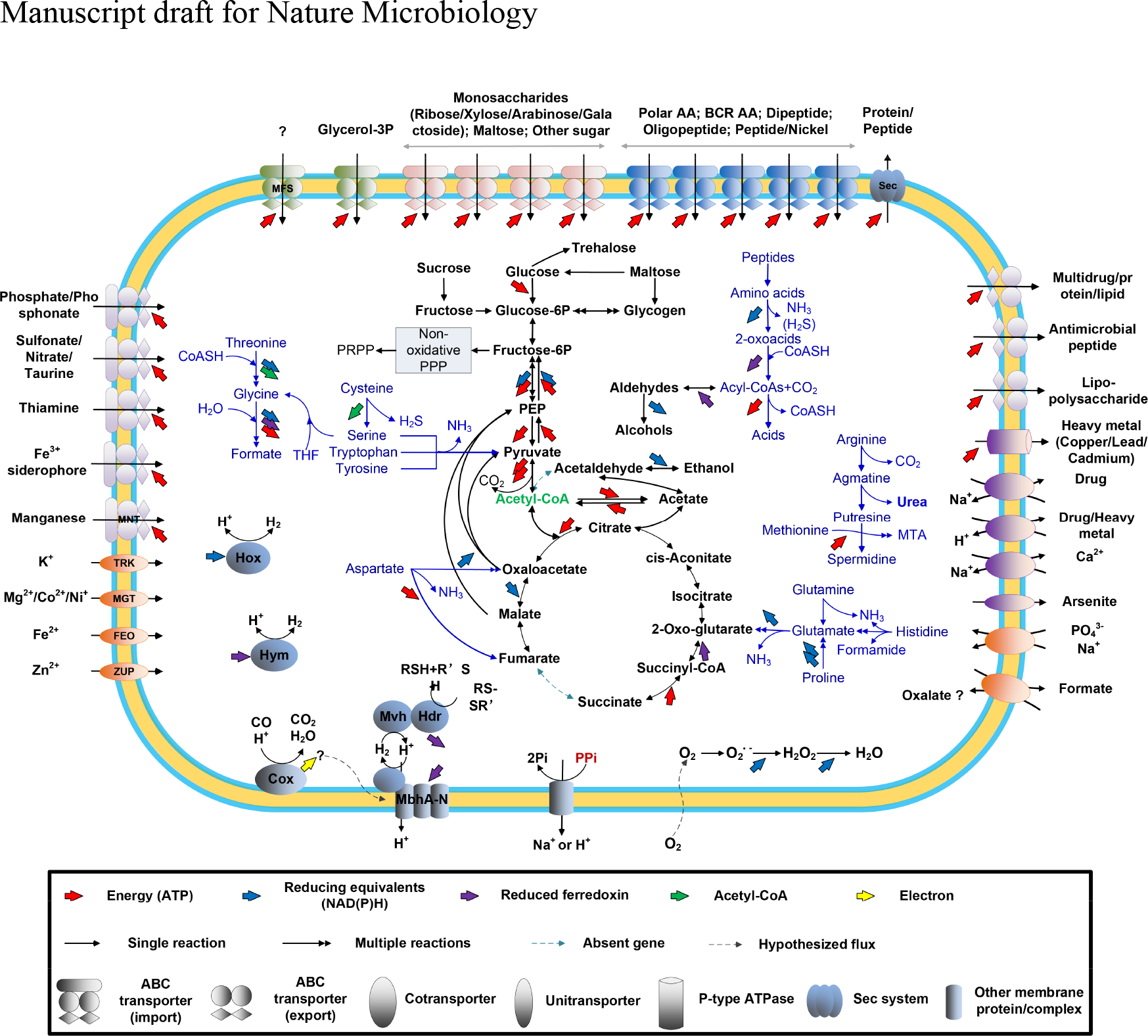
Metabolic model of Acetothermia sp. Ran1 based on the annotated genome sequences (**Table S6**). AA = Amino acids; BRC AA = Branched-chain amino acids; Sec = Secretion system; Glycerol-3P = Glycerol-3-phosphate; PPP = Pentose phosphate pathway; PRPP = 5-Phospho-alpha-D-ribose-1-diphosphate; ATP = Adenosine triphosphate; CoA = Coenzyme A; THF = Tetrahydrofolate; NAD(P)H = Nicotinamide adenine dinucleotide (phosphate) hydrogen; Pi = Phosphate; PPi = Pyrophosphate; MTA = 5′-S-Methyl-5′-thioadenosine; MNT = Manganese transporter; TRK = Potassium (K) transporter; MGT= Magnesium transporter; FED = Ferrous iron (Fe^2+^) transporter; ZUP = Zinc (Zn) transporter; MFS = Major facilitator superfamily transporter. More details on amino acids and electron transport metabolisms are shown in **Figure S10-12**.

#### Carbon uptake and central metabolism

Several ABC transporter genes were detected, including those for importing amino acids, peptides, glycerol-3-phosphate, maltose, ribose, and alpha-glucoside. This indicates that Ran1 can take up these compounds at the expense of ATP or the proton motive force (PMF) and use them as carbon and energy sources.

Sugars imported can be catabolized through the Embden-Meyerhof-Parnas pathway. The ATP produced during the transformation of hexoses to pyruvate can provide the cells with energy. Besides hexoses, Ran1 may utilize a broad range of pentoses, as it has all the genes of the non-oxidative pentose phosphate pathway^60^. Ran1 also encoded the complete pathway for glycogen metabolism and the gene encoding a trehalose synthase. Therefore, glycogen and trehalose may serve as carbon and energy storage, which can be utilized to mitigate fluctuations in substrate availability^61,62^. Two extracellular glycosylases were identified, including a cellulase and a glycoside hydrolase. This indicates that Ran1 has some limited extracellular saccharolytic activity and can hydrolyze polysaccharides from the feeding sludge into simpler sugars.

The pyruvate generated from sugars can be converted to acetyl-CoA by the pyruvate:ferredoxin oxidoreductase complex (*porABC*), generating reduced ferredoxin (Fdred). Acetyl-CoA can then enter the fermentation pathway catalyzed by two acetyl-CoA synthetases (*acsA* or *acdA*), resulting in the production of acetate and energy in the form of ATP.

The genome encoded an incomplete tricarboxylic acid (TCA) pathway, in which a succinate dehydrogenase (sdhABCD)/fumarate reductase (frdABCD) complex was not annotated. The partial pathway may serve as a source of biosynthetic precursors for anabolic pathways, as in methanogens and some other anaerobic bacteria^63,64^.

Amino acids and peptides, imported by ABC transporters, represent a potential source of carbon, nitrogen, energy, and building blocks of the cell. Indeed, it was found that the genome encoded genes for catabolizing at least 13 of the 22 amino acids (**Figure S10 and S11**). Serine, glycine, cysteine, aspartate, glutamate, glutamine, histidine, tyrosine, and tryptophan can be deaminated and converted into either pyruvate, oxaloacetate, or 2-oxoglutarate (**Figure S10**). These intermediates are then further oxidized by the pyruvate:ferredoxin oxidoreductase (*por*) or 2-ketoglutarate ferredoxin oxidoreductase (*kor*) to generate acetyl-CoA or succinyl-CoA, which can then be cleaved to yield acetate or succinate and energy in the form of ATP^65,66^. Glycine and serine can alternatively be degraded to formate through the glycine cleavage system and tetrahydrofolate pathway^42^, concomitant with the generation of ATP and reducing equivalents (in the form of NADH and Fdred). Some key enzymes involved in the catabolism of branched-chain amino acids were absent in the annotated genome (**Figure S11**). It is therefore only the non-branched-chained amino acids that can be used as energy source.

Limited capacity for amino acid synthesis was encoded in the genome (**Table S6**), indicating that some of the imported amino acids need to be directly used in anabolic pathways^67^.

Ran1 does not have the necessary genes for nitrogen fixation and ammonia import. Amino acids are thus predicted to be a major source of nitrogen, as NH3 is produced from deamination and assimilated via the glutamine/glutamate synthase pathway ^68^. The high dependence of exogenous amino acids and the fact that Ran1 only encode a single extracellular protease imply high dependence on the proteolytic action of other members of the microbial community.

#### Energy conservation and electron flow

Ran1 encodes an energy-conserving, membrane-bounded hydrogenase complex (Mbh AN) (Figure 5 and **S12**) which can translocate protons across the membrane while catalysing Fdred-driving H2 production^19,69^. It enables the cell to establish a PMF from Fdred^70^. The produced H2 and Fdox can be recycled by another complex formed by the electron-bifurcating heterodisulfide reductase (Hdr A-C) and the methyl viologen reducing hydrogenase (Mvh D,G,A)^69^. In addition, a bidirectional [NiFe] hydrogenase complex (Hox E,F,U,H,Y) and a putative [Fe] hydrogenase (Hym AB) were also encoded. These complexes catalyze the electron transfer between H+/H2 with NAD(P)H/NAD(P)+^71,72^ and Fdred/Fdox^73^, respectively. These bidirectional hydrogenases are hypothesized to function as electron valves, balancing reductants in the cell^72^. As part of the energy recycling system, the membrane-integral pyrophosphatase (HppA) can also translocate H^+^ or Na^+^ to generate PMF, using the energy produced from hydrolysis of pyrophosphate (PPi)^74^.

Surprisingly, the genome does not encode any conventional ATP synthases, which are often used to generate ATP at the expense of the established PMF^75^. Loss of functional ATP synthase has also been reported for other strictly anaerobic fermenters, such as *Clostridium acetobutylicum^16^* and *Clostridium perfringens^11^.* The energy stored in the PMF is therefore most likely used for active transport of substrates^78^.

#### Sulfur metabolism

Ran1 does not encode pathways for sulphate reduction. However, the complex formed by the electron-bifurcating heterodisulfide reductase (Hdr A-C) and the methyl viologen reducing hydrogenase (Mvh D,G,A) could function as a polysulfide/disulfide oxidoreductase (**Figure S12**), as proposed for other anaerobic bacteria^18,79^. Ran1 may therefore have a potential role in sulphur transformations in digesters.

#### Stress response

The genome possesses several genes typical of anaerobic bacteria, such as the oxygen-sensitive class III ribonucleoside triphosphate reductase, ferredoxin oxidoreductases, and radical S-adenosyl-methionine-dependent (SAM) proteins (**Table S6**). The oxygen-required class-I and oxygen-tolerant class-II ribonucleotide triphosphate reductases were not found. However, Ran1 encodes several proteins predicted to counter oxidative damage, including superoxide reductase, ruberythrin, thioredoxins, peroxidases, thioredoxin reductase, and glutaredoxins, which may allow it to survive under microaerobic conditions.

### Ecological significance and concluding remarks

This study presents the first detailed insight into the morphology, physiology, and ecology of a member of the candidate phylum Acetothermia (former OP1). The bacterium was stably present in several mesophilic sludge digesters during a period of several years and represents a novel genus that includes other previously detected 16S rRNA gene sequences of Acetothermia in anaerobic bioreactors. Members of this genus from digesters at different WWTPs belong to the same OTU and demonstrate the same morphology. These digesters, from two individual WWTPs, have no link between each other in terms of operation, seeding microbiome and feedstock. It may indicate low diversity of Acetothermia bacteria in anaerobic digester environments. The metabolic reconstruction suggested that it is an anaerobic, fermentative bacterium involved in acidogenesis, producing organic acids (such as acetate and formate) and hydrogen from the fermentation of peptides, amino acids, and simple sugars (maltose, sucrose). It might also use polysulfide as an alternative electron acceptor to produce hydrogen sulphide (H2S)-thus contributing to the turnover of sulfur and production of H_2_S.

The heterotrophic way of life predicted for Ran1 is similar to that of the oil reservoir-associated Acetothermia bacterium 64_32^12^, which affiliates to the same family level clade **(Figure 2A**). In contrast, the other more distantly related described member of the phylum, *Candidatus* Acetothermum autotrophicum, is predicted to have an acetogenic lifestyle by CO_2_ fixation^21^.

Interestingly, this Acetothermia bacterium demonstrated an unusual morphology composed of a central rod cell and long prosthecae protruding from both poles of the rod. This type of morphology is rarely observed for bacteria outside the Alphaproteobacteria^80^, and it is the first time prosthecae have been shown for a candidate phylum bacterium. The model organism for prosthecate bacterium, *C. crescentus*, has revealed important insight into development of cell morphologies^53^. As Ran1 is distantly related to all known prosthecate bacteria, it is likely that this bacterium can shed new light on the evolution of cell morphology. The long and flexible prosthecae greatly expand the surface area of the cell and provide increased access to nutrients under nutrient-limiting conditions. This is supported by their abundance being restricted to digesters with relatively low levels of phosphorus and other nutrients.

The genome generated in this study is one of few closed genomes for uncultured candidate phyla and importantly provides the foundation for future study on pathway expression of the lineage with metatranscriptomics and metaproteomics. The design of FISH probes for the genus will facilitate future *in situ* studies of the genus in other systems.

Phylogenetic analyses of the Ran1 genome classified it as a novel genus within the phylum. We suggest that the closed genome should serve as the type material for this genus ^81,82^ and propose the following taxonomic names for the novel genus and species:

#### *Bipolaricaulis* gen. nov

Etymology. L. pref. *bis bi*, twice; N.L. adj. *polaris* (from L. n. *polus*, a pole), pertaining to the poles of the rod-shaped cell; L. masc. n. *caulis*, stalk; N.L. masc. n. Bipolaricaulis, stalks at both poles.

#### *Bipolaricaulis anaerobia* gen. et sp. nov

Etymology. an.a.e.ro’bi.a. N. L. F. adj. Gr. pref. an not; Gr. N. aer air; Gr.n.bios life; N.L. adj. anaerobia, anaerobe, that can live in the absence of oxygen; referring to the respiratory metabolism of the organism.

## Acknowledgments

This study was supported by Innovation Fund Denmark (NomiGas, grant 1305-00018B) MiDAS, Obelske Family Foundation, Villum Foundation, and Aalborg University. The AFM equipment and its application by Hüsnü Aslan were partially funded by the Carlsberg Foundation. We thank Kirsten Nørgaard and Lisbet Adrian for providing samples and physico-chemical data from the plants. The LABGeM (CEA/IG/Genoscope & CNRS UMR8030) and the France Génomique National infrastructure (funded as part of Investissement d’avenir program managed by Agence Nationale pour la Recherche, contract ANR-10-INBS-09) are acknowledged for support within the MicroScope annotation platform.

## Contributions

L.-P.H., P.H.N., M.A. and M.S.D. designed the experiments. L.-P.H., M.S.D., R.H.K., and S.M.K. contributed to the genome construction. L.-P.H. and M.S.D. are responsible for genome annotation. S.J.M. designed the FISH probes and performed FISH. H.A. and R.M. conducted AFM analysis. W.E.Y.F. conducted Raman analysis. L.-P.H. contributed to sample collection and preparation, physico-chemical analysis, and amplicon sequencing. L.-P.H. and M.S.D. drafted the manuscript. All authors contributed to discussion and revision of the paper.

## References

1. Falkowski, P. G., Fenchel, T. & Delong, E. F. The microbial engines that drive earth’s biogeochemical cycles. Science 320, 1034–1039 (2008).

2. McIlroy, S. J. et al. MiDAS 2.0: an ecosystem-specific taxonomy and online database for the organisms of wastewater treatment systems expanded for anaerobic digester groups. Database 2017, 1–9 (2017).

3. Huttenhower, C. et al. Structure, function and diversity of the healthy human microbiome. Nature 486, 207–214 (2012).

4. Didelot, X., Walker, A. S., Peto, T. E., Crook, D. W. & Wilson, D. J. Within-host evolution of bacterial pathogens. Nat. Rev. Microbiol. 14, 150–162 (2016).

5. Caporaso, J. G. et al. Ultra-high-throughput microbial community analysis on the Illumina HiSeq and MiSeq platforms. ISME J. 6, 1621–1624 (2012).

6. Parks, D. H. et al. Recovery of nearly 8,000 metagenome-assembled genomes substantially expands the tree of life. Nat. Microbiol. 903, 1–10 (2017).

7. Kirkegaard, R. H. et al. The impact of immigration on microbial community composition in full-scale anaerobic digesters. Sci. Rep. 7, 9343 (2017).

8. Hugenholtz, P., Pitulle, C., Hershberger, K. L. & Pace, N. R. Novel division level bacterial diversity in a Yellowstone hot spring novel division level. J. Bacteriol. 180, 366–376 (1998).

9. Eloe-Fadrosh, E. A. et al. Global metagenomic survey reveals a new bacterial candidate phylum in geothermal springs. Nat. Commun. 7, 10476 (2016).

10. Solden, L., Lloyd, K. & Wrighton, K. The bright side of microbial dark matter: lessons learned from the uncultivated majority. Curr. Opin. Microbiol. 31, 217–226 (2016).

11. Mukherjee, S. et al. 1,003 reference genomes of bacterial and archaeal isolates expand coverage of the tree of life. Nat. Biotechnol. 35, 676–683 (2017).

12. Hu, P. et al. Genome-resolved metagenomic analysis reveals roles for candidate phyla and other microbial community members in biogeochemical transformations in oil reservoirs. MBio 7, 1–12 (2016).

13. Rinke, C. et al. Insights into the phylogeny and coding potential of microbial dark matter. Nature 499, 431–437 (2013).

14. Albertsen, M. et al. Genome sequences of rare, uncultured bacteria obtained by differential coverage binning of multiple metagenomes. Nat. Biotechnol. 31, 533–538 (2013).

15. Moitinho-Silva, L. et al. Integrated metabolism in sponge-microbe symbiosis revealed by genome-centered metatranscriptomics. ISME J. 11, 1–16 (2017).

16. Dick, G. J. & Baker, B. J. Omic approaches in microbial ecology: charting the unknown. Microbe 8, 353–360 (2013).

17. Vanwonterghem, I., Jensen, P. D., Ho, D. P., Batstone, D. J. & Tyson, G. W. Linking microbial community structure, interactions and function in anaerobic digesters using new molecular techniques. Curr. Opin. Biotechnol. 27, 55–64 (2014).

18. Kirkegaard, R. H. et al. Genomic insights into members of the candidate phylum Hyd24-12 common in mesophilic anaerobic digesters. ISME J. 10, 1–13 (2016).

19. Nobu, M. K. et al. Phylogeny and physiology of candidate phylum ‘Atribacteria’ (OP9/JS1) inferred from cultivation-independent genomics. ISME J. 10, 273–286 (2016).

20. McIlroy, S. J. et al. Genomic and in situ investigations of the novel uncultured Chloroflexi associated with 0092 morphotype filamentous bulking in activated sludge. ISME J. 10, 2223–2234 (2016).

21. Takami, H. et al. A deeply branching thermophilic bacterium with an ancient Acetyl-CoA pathway dominates a subsurface ecosystem. PLoS One 7, (2012).

22. Caporaso, J. G. et al. Global patterns of 16S rRNA diversity at a depth of millions of sequences per sample. Proc. Natl. Acad. Sci. 108, 4516–4522 (2011).

23. Albertsen, M., Karst, S. M., Ziegler, A. S., Kirkegaard, R. H. & Nielsen, P. H. Back to basics - the influence of DNA extraction and primer choice on phylogenetic analysis of activated sludge communities. PLoS One 10, e0132783 (2015).

24. Quast, C. et al. The SILVA ribosomal RNA gene database project: improved data processing and web-based tools. Nucleic Acids Res. 41, 590–596 (2013).

25. Bolger, A. M., Lohse, M. & Usadel, B. Trimmomatic: a flexible trimmer for Illumina sequence data. Bioinformatics 30, 2114–20 (2014).

26. Magoč, T. & Salzberg, S. L. FLASH: fast length adjustment of short reads to improve genome assemblies. Bioinformatics 27, 2957–2963 (2011).

27. Edgar, R. C. Search and clustering orders of magnitude faster than BLAST. Bioinformatics 26, 2460–2461 (2010).

28. Wang, Q., Garrity, G. M., Tiedje, J. M. & Cole, J. R. Naïve Bayesian classifier for rapid assignment of rRNA sequences into the new bacterial taxonomy. Appl. Environ. Microbiol. 73, 5261–5267 (2007).

29. R Core Team. R: a language and environment for statistical computing. R Foundation for Statistical Computing, Vienna, Austria. (2017). Available at: http://www.r-project.org/.

30. Wickham, H. ggplot2: elegant graphics for data analysis. Springer-Verlag New York (2009). Available at: http://ggplot2.org.

31. Boetzer, M. & Pirovano, W. SSPACE-LongRead: scaffolding bacterial draft genomes using long read sequence information. BMC Bioinformatics 15, 1–9 (2014).

32. Boetzer, M. & Pirovano, W. Toward almost closed genomes with GapFiller. Genome Biol. 13, R56 (2012).

33. Vallenet, D. et al. MicroScope: a platform for microbial genome annotation and comparative genomics. Database (Oxford). 2009, bap021 (2009).

34. Vallenet, D. et al. MicroScope--an integrated microbial resource for the curation and comparative analysis of genomic and metabolic data. Nucleic Acids Res. 41, D636–D647 (2013).

35. Ludwig, W. et al. ARB: a software environment for sequence data. Nucleic Acids Res. 32, 1363–1371 (2004).

36. Yilmaz, L. S., Parnerkar, S. & Noguera, D. R. MathFISH, a web tool that uses thermodynamics-based mathematical models for in silico evaluation of oligonucleotide probes for fluorescence in situ hybridization. Appl. Environ. Microbiol. 77, 1118–1122 (2011).

37. Cole, J. R. et al. Ribosomal Database Project: data and tools for high throughput rRNA analysis. Nucleic Acids Res. 42, D633–42 (2014).

38. McIlroy, S. J., Tillett, D., Petrovski, S. & Seviour, R. J. Non-target sites with single nucleotide insertions or deletions are frequently found in 16S rRNA sequences and can lead to false positives in fluorescence in situ hybridization (FISH). Env. Microbiol 13, 38–47 (2011).

39. Daims, H., Stoecker, K. & Wagner, M. Fluorescence in situ hybridization for the detection of prokaryotes. Mol. Microb. Ecol. 213, 239 (2005).

40. Greuter, D., Loy, A., Horn, M. & Rattei, T. probeBase—an online resource for rRNA-targeted oligonucleotide probes and primers: new features 2016. Nucleic Acids Res. 44, D586–D589 (2016).

41. Wallner, G., Amann, R. & Beisker, W. Optimizing fluorescent in situ hybridization with rRNA-targeted oligonucleotide probes for flow cytometric identification of microorganisms. Cytometry 14, 136–143 (1993).

42. Nobu, M. K. et al. Microbial dark matter ecogenomics reveals complex synergistic networks in a methanogenic bioreactor. ISME J. 9, 1710–1722 (2015).

43. Merhej, V., Royer-carenzi, M., Pontarotti, P. & Raoult, D. Massive comparative genomic analysis reveals convergent evolution of specialized bacteria. Biol. Direct 4, I3 (2009).

44. Perkins, S. D., Scalfone, N. B. & Angenent, L. T. Comparative 16S rRNA gene surveys of granular sludge from three upflow anaerobic bioreactors treating purified terephthalic acid (PTA) wastewater. Water Sci. Technol. 64, 1406–1412 (2011).

45. Kwon, S., Kim, T. S., Yu, G. H., Jung, J. H. & Park, H. D. Bacterial community composition and diversity of a full-scale integrated fixed-film activated sludge system as investigated by pyrosequencing. J. Microbiol. Biotechnol. 20, 1717–1723 (2010).

46. Goux, X. et al. Microbial community dynamics in replicate anaerobic digesters exposed sequentially to increasing organic loading rate, acidosis, and process recovery. Biotechnol. Biofuels 8, 122 (2015).

47. Chaganti, S. R., Lalman, J. A. & Heath, D. D. 16S rRNA gene based analysis of the microbial diversity and hydrogen production in three mixed anaerobic cultures. Int. J. Hydrogen Energy 37, 9002–9017 (2012).

48. Yarza, P. et al. Uniting the classification of cultured and uncultured bacteria and archaea using 16S rRNA gene sequences. Nat. Rev. Microbiol. 12, 635–645 (2014).

49. Kysela, D. T., Randich, A. M., Caccamo, P. D. & Brun, Y. V. Diversity takes shape: understanding the mechanistic and adaptive basis of bacterial morphology. PLoS Biol. 14, e1002565 (2016).

50. Porter, J. S. & Pate, J. L. Prosthecae of Asticcacaulis biprosthecum: system for the study of membrane transport. J. Bacteriol. 122, 976–986 (1975).

51. Wagner, J. K., Setayeshgar, S., Sharon, L. A., Reilly, J. P. & Brun, Y. V. A nutrient uptake role for bacterial cell envelope extensions. Proc. Nat. Acad. Sci. U. S. A. 103, 11772–11777 (2006).

52. McAdams, H. H. Bacterial stalks are nutrient-scavenging antennas. Proc. Nat. Acad. Sci. U. S. A 103, 11435–11436 (2006).

53. Woldemeskel, S. A. & Goley, E. D. Shapeshifting to survive: shape determination and regulation in Caulobacter crescentus. Trends Microbiol. 25, 673–687 (2017).

54. Vasilyeva, L. V et al. Asticcacaulis benevestitus sp. nov., a psychrotolerant, dimorphic, prosthecate bacterium from tundra wetland soil. Int. J. Syst. Evol. Microbiol. 56, 2083–2088 (2017).

55. Gonin, M., Quardokus, E. M., Donnol, D. O. & Maddock, J. Regulation of stalk elongation by phosphate in Caulobacter crescentus. J. Bacteriol. 182, 337–347 (2000).

56. Sutcliffe, I. C. A phylum level perspective on bacterial cell envelope architecture. Trends Microbiol. 18, 464–470 (2010).

57. Huber, R. et al. Thermotoga maritima sp. nov. represents a new genus of unique extremely thermophilic eubacteria growing up to 90°C. Arch. Microbiol. 144, 324–333 (1986).

58. Jiang, Y., Zhou, Q., Wu, K., Li, X. Q. & Shao, W. L. A highly efficient method for liquid and solid cultivation of the anaerobic hyperthermophilic eubacterium Thermotoga maritima. FEMS Microbiol. Lett. 259, 254–259 (2006).

59. Imam, S., Chen, Z., Roos, D. S. & Pohlschröder, M. Identification of surprisingly diverse type IV pili, across a broad range of gram-positive bacteria. PLoS One 6, e28919 (2011).

60. Stincone, A. et al. The return of metabolism: biochemistry and physiology of the pentose phosphate pathway. Biol. Rev. Camb. Philos. Soc. 90, 927–963 (2014).

61. Wilson, W. A. et al. Regulation of glycogen metabolism in yeast and bacteria. FEMS Microbiol. Rev. 34, 952–985 (2010).

62. Arguelles, J. C. Physiological roles of trehalose in bacteria and yeasts: a comparative analysis. Arch. Microbiol. 174, 217–224 (2000).

63. Rosenberg, E., DeLong, E. F., Lory, S., Stackebrandt, E. & Thompson, F. The Prokaryotes: prokaryotic physiology and biochemistry. (Springer-Verlag Berlin Heidelberg, 2013). doi:10.1007/978-3-642-30141-4

64. Nobu, M. K., Narihiro, T., Kuroda, K., Mei, R. & Liu, W. T. Chasing the elusive Euryarchaeota class WSA2: genomes reveal a uniquely fastidious methyl-reducing methanogen. ISME J. 2, 1–10 (2016).

65. Adams, M. W. W. et al. Key role for sulfur in peptide metabolism and in regulation of three hydrogenases in the hyperthermophilic archaeon Pyrococcus furiosus. J. Bacteriol. 183, 716–724 (2001).

66. Fukui, T. et al. Complete genome sequence of the hyperthermophilic archaeon Thermococcus kodakaraensis KOD1 and comparison with Pyrococcus genomes. Genome Res. 15, 352–363 (2005).

67. Brown, C. T. et al. Unusual biology across a group comprising more than 15% of domain Bacteria. Nature 523, 208–211 (2015).

68. Bravo, A. & Mora, J. Ammonium assimilation in Rhizobium phaseoli by the glutamine synthetase-glutamate synthase pathway. J. Bacteriol. 170, 980–984 (1988).

69. Buckel, W. & Thauer, R. K. Energy conservation via electron bifurcating ferredoxin reduction and proton/Na+ translocating ferredoxin oxidation. Biochim. Biophys. Acta-Bioenerg. 1827, 94–113 (2013).

70. Sapra, R., Bagramyan, K. & Adams, M. W. W. A simple energy-conserving system: proton reduction coupled to proton translocation. Proc. Nat. Acad. Sci. U. S. A 100, 7545–7550 (2003).

71. Cassier-Chauvat, C., Veaudor, T. & Chauvat, F. Advances in the function and regulation of hydrogenase in the cyanobacterium Synechocystis PCC6803. Int. J. Mol. Sci. 15, 19938–19951 (2014).

72. Eckert, C. et al. Genetic analysis of the Hox hydrogenase in the cyanobacterium Synechocystis sp. PCC 6803 reveals subunit roles in association, assembly, maturation, and function. J. Biol. Chem. 287, 43502–43515 (2012).

73. Fritsch, J., Lenz, O. & Friedrich, B. Structure, function and biosynthesis of O_2_-tolerant hydrogenases. Nat. Rev. Microbiol. 11, 106–14 (2013).

74. Luoto, H. H., Baykov, A. a, Lahti, R. & Malinen, A. M. Membrane-integral pyrophosphatase subfamily capable of translocating both Na^+^ and H^+^. Proc. Nat. Acad. Sci. U. S. A 110, 1255–60 (2013).

75. Mayer, F. & Mu, V. Adaptations of anaerobic archaea to life under extreme energy limitation. FEMS Microbiol. Rev. 38, 449–472 (2014).

76. Nolling, J. et al. Genome sequence and comparative analysis of the solvent-producing bacterium Clostridium acetobutylicum. J. Bacteriol. 183, 4823–4838 (2001).

77. Shimizu, T. et al. Complete genome sequence of Clostridium perfringens, an anaerobic flesh-eater. Proc. Nat. Acad. Sci. U. S. A 99, 996–1001 (2002).

78. Yan, N. Structural investigation of the proton-coupled secondary transporters. Curr. Opin. Struct. Biol. 23, 483–491 (2013).

79. Nobu, M. K. et al. The genome of Syntrophorhabdus aromaticivorans strain UI provides new insights for syntrophic aromatic compound metabolism and electron flow. Environ. Microbiol. 17, 4861–4872 (2015).

80. Randich, A. M. & Brun, Y. V. Molecular mechanisms for the evolution of bacterial morphologies and growth modes. 6, 1–13 (2015).

81. Whitman, W. B. Genome sequences as the type material for taxonomic descriptions of prokaryotes. Syst. Appl. Microbiol. 38, 217–222 (2015).

82. Whitman, W. B. Modest proposals to expand the type material for naming of prokaryotes. Int. J. Syst. Evol. Microbiol. 66, 2108–2112 (2016).

83. Parks, D. H., Imelfort, M., Skennerton, C. T., Hugenholtz, P. & Tyson, G. W. CheckM: assessing the quality of microbial genomes recovered from isolates, single cells, and metagenomes. Genome Res. 25, 1043–55 (2015).

